# Atf4 protects islet β-cell identity and function under acute glucose-induced stress but promotes β-cell failure in the presence of free fatty acid

**DOI:** 10.1101/2024.06.28.601249

**Authors:** Mahircan Yagan, Sadia Najam, Ruiying Hu, Yu Wang, Prasanna Dadi, Yanwen Xu, Alan J. Simmons, Roland Stein, Christopher M. Adams, David A. Jacobson, Ken Lau, Qi Liu, Guoqiang Gu

**Affiliations:** Department of Cell and Developmental Biology, Vanderbilt University School of Medicine, Nashville, TN 37232, USA; Center for Stem Cell Biology, Vanderbilt University School of Medicine, Nashville, TN 37232, USA; Department of Biostatistics and Center for Quantitative Sciences, Vanderbilt Medical Center, Nashville, TN37232, USA; Department of Molecular Physiology and Biophysics, Vanderbilt University School of Medicine, Nashville, TN 37232, USA; Epithelial Biology Center, Vanderbilt Medical Center, Nashville, TN 37232, USA; Division of Endocrinology, Diabetes, Metabolism and Nutrition. Mayo Clinic, Rochester, MN55905.

## Abstract

Glucolipotoxicity, caused by combined hyperglycemia and hyperlipidemia, results in β-cell failure and type 2 diabetes (T2D) via cellular stress-related mechanisms. Activating transcription factor 4 (Atf4) is an essential effector of stress response. We show here that *Atf4* expression in β-cells is dispensable for glucose homeostasis in young mice, but it is required for β-cell function during aging and under obesity-related metabolic stress. Henceforth, aged *Atf4-*deficient β-cells display compromised secretory function under acute hyperglycemia. In contrast, they are resistant to acute free fatty acid-induced loss-of identity and dysfunction. At molecular level, *Atf4*-deficient β-cells down-regulate genes involved in protein translation, reducing β-cell identity gene products under high glucose. They also upregulate several genes involved in lipid metabolism or signaling, likely contributing to their resistance to free fatty acid-induced dysfunction. These results suggest that *Atf4* activation is required for β-cell identity and function under high glucose, but this paradoxically induces β-cell failure in the presence of high levels of free fatty acids. Different branches of Atf4 activity could be manipulated for protecting β-cells from metabolic stress-induced failure.

**Highlights:**

- Atf4 is dispensable in β-cells in young mice
- Atf4 protects β-cells under high glucose
- Atf4 exacerbate fatty acid-induced β-cell defects
- Atf4 activates translation but depresses lipid-metabolism

## Introduction

Type 2 diabetes (T2D) usually starts with obesity-induced insulin resistance, which demands higher insulin output from islet β-cells for glucose homeostasis. Yet high insulin secretion, together with the high levels of glucose and free fatty acid metabolism, causes β-cell dysfunction, loss of identity, and/or death (i.e. β-cell failure) (1,2), followed by overt T2D. Thus, finding ways to delay or to reverse β-cell failure can delay or cure T2D.

A major inducer of β-cell failure is cellular stress (3). For high levels of insulin secretion, β-cells metabolize a large flux of glucose and handle a large load of insulin biosynthesis/processing/folding. These events generate reactive oxygen species (ROS) and unfolded proteins (4,5), stressors that at high levels reduce the ER function and/or the levels/activities of key β-cell factors (6). Similarly, free fatty acids can acutely induce insulin secretion via G protein-coupled receptors and intracellular metabolism (7-9). They also induce the production of ceramide and ROS, leading to cellular stress and ER/mitochondrial dysfunction (10,11). Therefore, β cells constantly activate the oxidative stress response and unfolded-protein response (UPR) for homeostasis, especially under obesity-related hyperglycemia and hyperlipidemia (i.e., glucolipotoxicity) (12,13). Intriguingly, stress response can be overactivated, leading to β-cell failure due to limited translation of key β-cell factors and/or activation of cell death (14,15). Thus, both failed and overactivated stress response have been associated with human diabetes (16,17). The function of Atf4 (activating transcription factor 4) is particularly relevant to the double-edged nature of stress response. It is activated during integrated stress response by several signals including ER stress (18). It can activate or repress the transcription of its target, likely via binding to different co-regulators (19). Intriguingly, its short-term activation leads to homeostasis by inducing factors involved in lipid/glucose metabolism and homeostasis (20-23); but its overactivation induces C/EBP homologous protein (CHOP, encoded by *DDIT3*), causing cell death (24,25) (26).

Several studies have attempted to address the function of *Atf4* in islet β-cells. Specifically, Atf4 targets or interacting proteins, e.g., Eif4bp1 and TRB3, have been shown to regulate β-cell function and survival (23,27). Recently, we reported that *Atf4* overexpression in mouse β-cells compromises their identity and function (15). Two other studies reported the roles of *Atf4* in β-cell function and viability under induced ER stress and normal physiology (22,28). However, these latter studies inactivated *Atf4* using a *Rip^Cre-TG^* transgenic line (22,28), which was reported to compromise islet function via ectopic expression of human growth hormone (29). In addition, the β-cell specific gene expression in *Atf4* mutants was not analyzed, leaving a gap in understanding how Atf4 regulates β-cell function. Here, we report our findings utilizing both *Rip^Cre-TG^* and the *Ins1^Cre^* knockin line to inactivate *Atf4* in mouse β-cells. We uncover the opposing roles of *Atf4* in β-cells under acute hyperglycemia or hyperlipidemia conditions.

## Research Design and Methods

### Animal models and procedures

Mouse usage followed protocol approved by the Vanderbilt IACUC for Dr. Gu, in compliance with the policies of AALAC. C57Bl/6 mice, *MT/MG* [*Gt(ROSA)26Sor^tm4(ACTB-tdTomato,-EGFP)Luo^*/J] mice, *Rosa26^eYFP^* [*Gt(ROSA)26Sor^tm1(EYFP)Cos^*/J] mice, *Rip^Cre-TG^* (B6.Cg-Tg(Ins2-Cre)25Mgn/J), *Ins1^Cre^* [B6(Cg)-*Ins1^tm1.1(cre)Thor^*/J] were from Jackson Laboratory. *Atf4^F/F^* mice were reported (30). S961 infusion (gift from Nova Nordisk) uses an alzet osmotic pump (#2001). Note that mice used are mostly CBA/Bl6 background (>95%), estimated based on crossing history.

### Consideration of sex in mouse models

Both male and female mice at equal ratio (when possible) are included. Data were presented separately or in combination as indicated in each figure.

### β cell preparation, secretion, and ATP/ADP assays

Islet isolation uses collagenase Type IV digestion, with postnatal day 4 (P4) pancreata directly digested while older mice via infusion (31). For secretion assays, islets were allowed to recover in complete RPMI1066 media (with antibiotics and 10% FBS) for overnight. They were directly used for secretion assays or further treated for ∼48 hours with 5.5 mM glucose (G5.5), G20, or (G5.5 + 0.6mM palmitate) (Pa0.6) followed by secretion assays. Insulin secretion follow standard protocols (31), induced with G2.8, G20, or (G20 + 30mM KCl) (G20K). For each genotype/condition, two-three biological replications with at least three technical replications for each were included. Total insulin was assayed using alcohol-acid extraction. Insulin was measured with an Elisa kit from ALPCO. The % of total insulin secreted within a 45-minute window was presented. ADP/ATP ratio assays use an EnzyLight^TM^ ADP/ATP ratio Assay Kit (BioAssay Syetems).

### Intraperitoneal glucose tolerance test (IPGTT) and high-fat diet (HFD)

IPGTT followed previously described method (32). Mice were fasted overnight. Glucose was injected at 2mg/kg. Blood glucose was then measured via tail nip. To induce obesity, 5-6 weeks-old mice were fed with HFD for three months. The high fat diet is from VWR, with 60% calories from fat (compare with 3% fat in control diet).

### IF assays in tissue sections or whole islets

IF assays follow routine method using cells spun onto slides, whole islets, both frozen and paraffin sections. To assay for β-cell proliferation, islets were dissociated into single cells and spun onto slides. They were stained for Ki67, insulin, and glucagon. For protein level-per-cell assays, whole mount IF was used. For cell death assays and counting islet cell types, tissue sections were used. After staining, images were taken with an Olympus FV-1000 confocal microscope. Cells with desired marker labeling were then identified (e.g, Ki67, Ins+ or Gcg+ cells) and counted.

Antibodies used are: Guinea pig anti-insulin (Dako), Goat anti-insulin (Jackson ImmunoResearch), Rabbit anti-MafA (Novis), guinea pig anti-Pdx1 (gift from Wright), goat anti-SS (Jackson ImmunoResearch), mouse anti-Gcg (ABCam), and Rabbit anti-Ki67 (ABCam). Secondary antibodies are: Alexa-Flour-488-donkey anti-guinea pig, anti-goat, anti-rabbit, Alexa-Flour-568-donkey anti-guinea pig, anti-rabbit, anti-mouse, and Alexa-Flour-647-donkey anti-guinea pig, anti-rabbit are all from Jackson Immunoresearch or ABCam. All antibodies use 1:1000 dilution.

### Ca^2+^ recording

Ca^2+^ recording followed methods in (33). Islets were cultured on ploy-lysine coated glass for 48 hours in RPMI1066 medium + 10% FBS (G11). FURA2AM (Invitrogen) was then added to 2 mM for 25 minutes followed by 20 min in recording buffer. Islets were then transferred to REC buffer with G2.0 and recorded for 5-minutes followed by G20 for 20 minutes. The recorded are the ratios of emitted fluorescence at excitation of 340 and 380 nm (F340/F380) at 5-seconds intervals.

### ScRNA-seq and analysis

ScRNA-seq follow that in (34-36). Dissociated islets with ∼70-80% single cells and <5% dead cells were flowed into a microfluidic chip for droplet-formation, followed by reverse transcription, library preparation and sequencing (targeting 120 million reads. Novaseq 6000, Illumina). Each sample has islets from four mice (two males and two females). DropEst (37) was used to preprocess scRNA-seq reads and to generate count matrices. Cells with low uniquely mapping reads (<500), low proportion of expressed genes (<100) or high proportion mitochondrial RNAs (>10%) will be excluded (Fig. S3). Reads were normalized using UMI-filtered counts. Cell subpopulations were identified and visualized by UMAP using Seurat based on the first 30 principal components generated from the top 2000 highly variable genes (38,39). Cell subpopulations were automatically annotated by scMRMA (40), and then further manually checked by known marker genes. Differentially expressed genes (DEGs) between mutant and control β-cells were identified by Seurat at the criteria of |log2 fold change|> 0.25 & FDR< 0.05.

#### Bioinformatic gene set enrichment analysis

After identification of DEGs, the Database for Annotation, Visualization and Integrated Discovery (DAVID) was primarily utilized for comprehensive functional clustering analysis (41). In all cases, similar pathways or processes were consolidated and summarized for presentation. Only top 10 were presented. We also utilized the pre-ranked DEGs in GO term enrichment and GSEA utilizing the GSEApy package (40). The findings were presented in Table S2.

### Statistical analysis

Statistical analyses utilized standard Student’s *t*-test for pairwise comparisons at single time points or paired genotypes. Two-way ANOVA were for comparing multiple groups of data points. Hypergeometric analyses were used for assessing the level of enrichment between list of genes. A *p*-value of 0.05 or lower was considered significant.

### Data Accessibility and request of materials

The RNAseq data will be available in the Gene Expression Omnibus (GEO) database. Further requests for resources and reagents should be directed to and will be fulfilled by the Lead Contact (615-936-3634).

## Results

### Establishing β-cell specific *Atf4* inactivation

To examine the function of *Atf4* in islet β cells, we initially inactivated this gene utilizing *Rip^Cre-TG^* mice as reported in (22). Unlike reported, our *Atf4^F/F^; Rip^Cre-TG^* mice have similar glucose clearance compared to *Atf4^F/+^; Rip^Cre-tg^* and *Rip^Cre-tg^*controls as late as 6 months of age (Fig. S1A). Corresponding to these results, active Cre was detected in only a portion of β-cells (Fig. S1B). We therefore switched to *Ins1^Cre^* for *Atf4* inactivation. *Ins1^Cre^*expresses active Cre in nearly all β-cells (Fig. S1C) (43). There is no detectable difference in the glucose tolerance of *Atf4^F/F^*and *Atf4^F/+^*; *Ins1^Cre^* or *Ins1^Cre^* mice. Neither glucose-stimulated insulin secretion (GSIS, induced by 20 mM glucose or G20) or KCl-stimulated insulin secretion (KSIS, induced by 30 mM KCl plus G20, or G20K) from their islets (Fig. S1D, E). *Atf4^F/F^*littermates were therefore used as controls for all below studies.

### Atf4 is required for postnatal β-cell proliferation

We examined if *Atf4* is required for β-cell production based on the roles of stress response in the process (44,45). At one-month of age, there is no difference in the numbers and sizes of islets that can be recovered from each control (*Atf4^F/F^*) or mutant (*Atf4^F/F^; Ins1^Cre^*) pancreas (Fig. S2A-C). The α-to-β cell ratio is higher in mutant islets (Fig. 1A-C), in both male and female mice. We did not detect altered δ-to-β cell ratio (Fig. 1C). The increased α-to-β cell ratio developed after birth, because no difference was detected at P4 (postnatal day 4) (Fig. 1D-F). Intriguingly, mutant- and control β-cells have similar level of proliferation (Fig. 1D-F), but the α-cells in mutants proliferate more for unknown reasons (Fig. 1F), likely contributing to the increased α-to-β ratio. In the rest of the studies, we focused on how *Atf4* inactivation in β-cells impacts their function and mouse physiology.

**Fig. 1.**
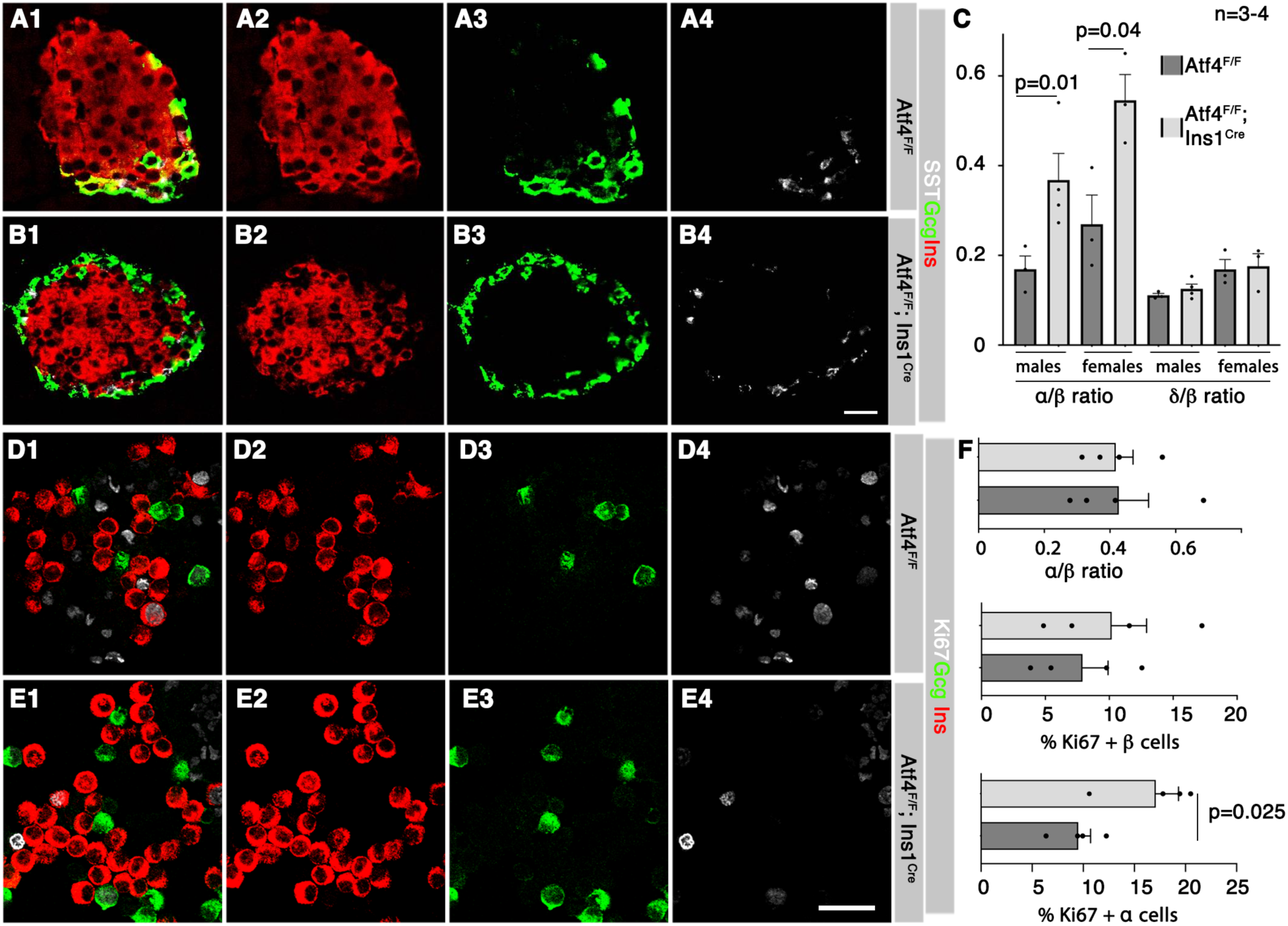
Atf4-inactivation in β-cells results in decreased β-to-α cell ratio in adolescent mice. For ease of tracking, panels labeled with a same capital letter are different channels of a single tissue section. (A-C) Ins, Gcg, and Sst IF staining and quantification to examine the ratio of between these islet cell types in one-month old mice control (*Atf4^F/F^*) and mutant (*Atf4^F/F^; Ins1^cre^*) islets. The quantification data in C are mean + SEM (n=3-4, males and females are shown separately). P-values are from unpaired, 2-tailed, type 2 student t-test. (D-F) IF staining and quantification of Ins, Gcg, and Ki67 in P4 islet cells, cyto-spun onto slides. The quantification data in F are mean + SEM (n=4). Two males and two females are included and presented together. P-values are from unpaired, 2-tailed, type 2 student t-test. Scale bars = 20 μm.

### Atf4 is dispensable for postnatal β-cell function and survival under normal physiology, but it is required under metabolic stress

Both male and female *Atf4^F/F^*; *Ins1^Cre^* mice have similar glucose clearance as *Atf4^F/F^* controls 1- and 4-months after birth (Fig. 2A-F). So are their GSIS and KSIS from isolated islets (Fig. S2D, E and below Fig. 3D). These findings suggest that *Atf4* is largely dispensable for endocrine function in β cells under normal physiology, in contrast to that reported in (22). For unknown reasons, the one-month-old *Atf4^F/F^*; *Ins1^Cre^* male mice have lower body weight than controls (Fig. 2C). Yet their weight caught up latter (Fig. 2F) and this property was not followed up.

**Fig. 2.**
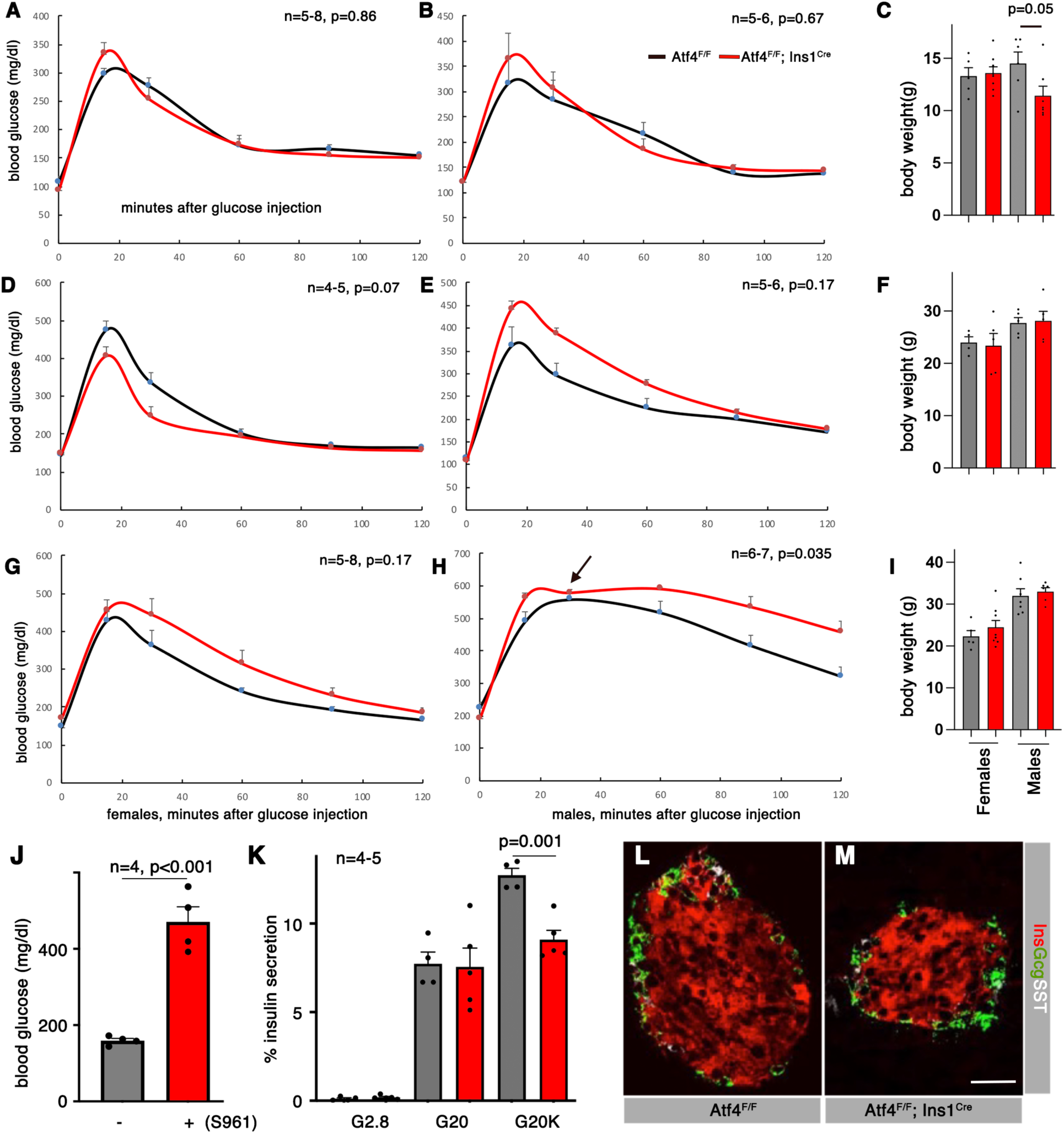
*Atf4* is dispensable for glucose homeostasis under normal feeding condition but is required under high-fat-diet challenge. For glucose homeostasis tests under normal and obesity conditions, mice of different age were used for IPGTT (A-I). For testing the effects of acute hyperglycemia, S961, via minipump-delivery, was used in two-months old mice (J-M). (A-C) IPGTT results from one-month old mice, with male and female data presented separately. Presented are mean + SEM. P value is calculated with unpaired, 2-tailed, type 2 student t-test. (D-F) IPGTT results as (A-C) except that 4-months old mice were used. (G-I) IPGTT results as (A-C) except mice were fed with HFD for three months. Arrow in H indicates a time point when several mutant mice have readings >600 mg/dl (the limit of glucometer). Their reading was arbitrarily set at 600, likely contributing to the depressed average at this point. (J) Random blood glucose one day after S961 infusion. (mean + SEM) are shown. P is from unpaired, 2-tailed, type 2 student t-test. (K) Insulin secretion from islets isolated from control and mutant mice (both male and females included) that have received S961 infusion for seven days. Mean + SEM were shown, p value is from unpaired, 2-tailed, type 2 student t-test. (L, M) Representative islet morphology after control and mutant male mice that have received S961 infusion for 7 days. Scale bars = 20 μm.

**Fig. 3.**
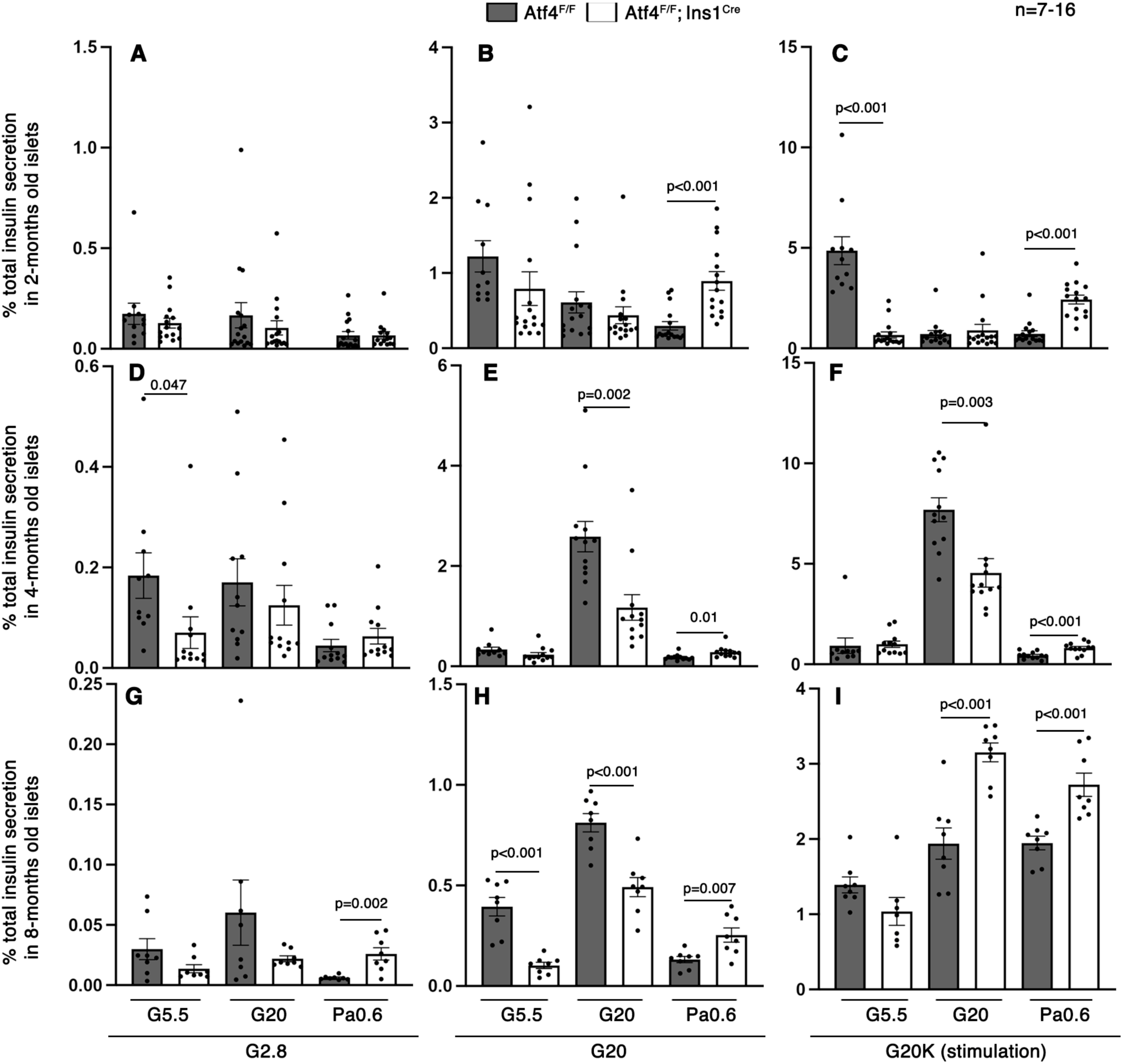
Atf4-deficient β-cells are liable to high-glucose induced dysfunction but are resistant to free fatty acid-induced dysfunction. Islets were isolated from 2-, 4-, and 8-months mice. They were then treated with G5.5, G20, or Pa0.6 for 48 hours, followed by insulin secretion assays. Shown are % of total insulin secretion within 45-minutes windows. (A-C) Results from 2-months-old islets. (D-F). Results from 4-months old mice. (G-I). Eight-months-old mice. (A, D, G) Basal insulin secretion induced by G2.8. (B, E, H) GSIS induced by G20. (C, F, I) KSIS induced by high glucose and KCl (G20K). For all panels, presented are mean + SEM. The *p* values were calculated with unpaired, 2-tailed, type 2 student t-test.

We next tested whether *Atf4-*deficient β cells are more sensitive to obesity-induced failure. After three-months of HFD treatment, female mutant mice displayed a weak trend of glucose intolerance (Fig. 2G); while male mutants showed a significantly compromised tolerance despite their same levels of weight gain when compared with controls (Fig. 2H, I). These sex-dependent phenotypes are consistent with the findings that male rodents are more liable for diabetes development (46).

We then tested the response of the *Atf4^F/F^*; *Ins1^Cre^* β cells to acute hyperglycemia. Acute insulin resistance was induced using S961, resulting in hyperglycemia within 24 hours (Fig. 2J). A week after, islets from S961-treated *Atf4^F/F^*; *Ins1^Cre^* mice have similar levels of basal insulin secretion (stimulated with G2.8) and GSIS as controls (Fig. 2K). Yet the mutant islets have significantly lowered KSIS (Fig. 2K). We did not notice altered islet morphology (i.e., with Ins^+^ cells locate in islet core while other islet cell types in periphery (Fig. 2L, M). Neither did we observe cell death in islets, examined via TUNEL assays (Fig. S2D-E).

These above findings suggest that *Atf4* is required for maintaining the pool of releasable insulin but not for β-cell viability under acute hyperglycemia. We therefore examined how *Atf4*-deficiency affects secretion, focusing on the effects of aging or obesity-related factors (hyperglycemia and hyperlipidemia).

### *Atf4* regulates insulin secretion in age- and stress type-specific manners

Islets were isolated from 2-, 4-, and 8-months old control and *Atf4^F/F^*; *Ins1^Cre^* mutant mice. They were treated for 48 hours in culture with G5.5, G20, and 0.6 mM palmitate together with G5.5 (Pa0.6), respectively. They were then examined for insulin secretion. For initial analysis, we processed the data from 2- and 4-months old male and female islets separately and observed similar results (Fig. S2F). We therefore presented combined data from both sexes from now on.

For 2-months-old samples, the basal insulin secretion levels in mutant islets are the same compared to their controls, pre-treated by G5.5, G20, or Pa0.6 (Fig. 3A). For GSIS, Pa0.6 treated mutant islets secrete more than controls (Fig. 3B). For KSIS, G5.5-treated mutant islets secrete less while Pa0.6-treated mutant islets secrete more (Fig. 3C). These results suggest that Atf4 increases the amount of releasable insulin at low glucose, but it represses in the presence of free fatty acid.

Compared with 2-months, 4- and 8-months old mutant islets display more pronounced difference compared with controls. Specifically, 4-months old islets have slightly reduced basal secretion under G5.5 treatment (Fig. 3D). Intriguingly, GSIS is lower but KSIS is higher in G20-treated mutant islets compared with controls (Fig. 3E). Same results were observed for KSIS at this stage (Fig. 3F). Along a similar line, eight-months old mutant islets developed additional secretory defects. Significantly, the basal secretion is higher in Pa0.6-treated mutant islets (Fig. 3G). GSIS is lower in both G5.5- and G20-treated mutant islets but higher in Pa0.6-treated mutant islets (Fig. 3H). For KSIS, both G20-treated and Pa0.6-treated mutant islets secrete more than controls (Fig. 3I).

These combined data underscore a complicated Atf4-regulated response, with both aging and stress-type as key determinant. Yet a basic theme is that Atf4 promotes β-cell GSIS under acute high glucose but reduces it under free fatty acid. This conclusion is surprising because hyperlipidemia was known to synergize with hyperglycemia to induce β-cell dysfunction. We therefore asked if the total insulin per β-cell was decreased by Pa0.6 treatment, which would have artificially increased the % of total insulin secretion.

### *Atf4-*deficient β-cells produce less insulin under acute high glucose treatment but more in the presence of high levels of free fatty acids

Islets were treated with G5.5, G20, or Pa0.6 for 48 hours. They were then assayed for insulin, MafA, and Pdx1 levels, a few representative markers for β-cell identity and function. There is no significant difference in insulin levels of control and mutant β-cells when treated with G5.5 (Fig. 4A, B, G). Insulin level is reduced in mutant β-cells under G20 (Fig. 4C, D, G), but increased under Pa0.6 (Fig. 4E-G).

**Fig. 4.**
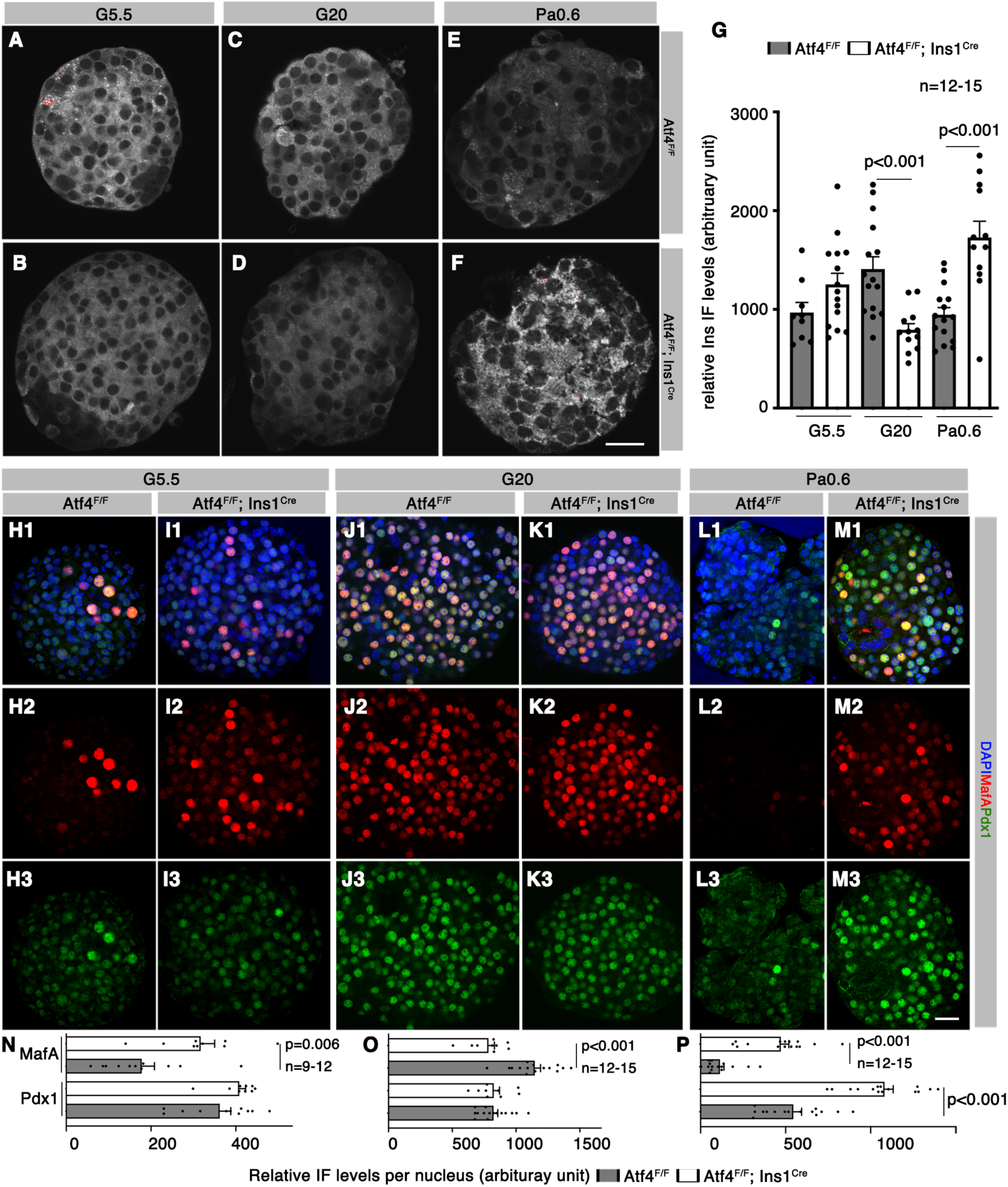
Atf4 promotes the expression of β-cell markers under high glucose but represses in the presence of free fatty acid. Four-months old islets were isolated and then treated with G5.5, G20, or Pa0.6 for 48 hours. They were then stained as whole mount to compare the expression of insulin, MafA, and Pdx1. Note that if different channels of a same section were presented, they will be labeled with a same capital letter, followed by a number to denote different antigens tested. (A-G) Insulin levels in islets treated G5.5, G20, and Pa0.6. Presented in G are mean + SEM. Note that in G, each dot represents a single islet section, including 40-200 cells each. The *p* values were calculated with unpaired, 2-tailed, type 2 student t-test. (H-P) Quantification of MafA and Pdx1 expression in treated islets. Note that in this case, Pdx1 and MafA were assayed using identical sections, presented as single and merged channels. In the merged channels, DAPI was included to locate the nuclei. The quantification data in N-P are mean +SEM, representing the levels at G5.5, G20, and Pa0.6, respectively. Each dot represents one islet section, including 66-179 nuclei. The *p* values were calculated with unpaired, 2-tailed, type 2 student t-test. Scale bars=20 μm.

Under G5.5 and G20 treatment, there is no difference in Pdx1 levels between control and mutant β-cells (Fig. 4H, J, N, O). Yet the MafA levels are higher in mutant β-cells under G5.5, but lower under G20 (Fig. 4I, K, N, O). Pa0.6 treatment induces higher Pdx1 and MafA levels in mutant β-cells (Fig. 4L, M, P). These findings, combined with the lowered insulin secretion in G20- but increased secretion in Pa0.6-treated mutant islets, suggest that Atf4 normally represses the production of releasable insulin under hyperlipidemia condition, while increase it under hyperglycemia. We next examined the Atf4-depenent gene expression in β-cells to establish the molecular basis of these functions.

### Atf4 regulates protein translation in β-cells

ScRNA-seq was utilized to identity the DEGs between control and *Atf4^F/F^*; *Ins1^Cre^* mutant β-cells at 2-months of age before any detectable defect appears. From two duplicated biological samples, we identified 6,394 high quality single-cell transcriptomes, with 17,389 genes with detectable expression (Fig. S3). The four endocrine islet cell types, enteroendocrine cells, endothelial cells, and macrophages were readily separable (Fig. 5A). Consistent with the β-cell specificity of *Ins1^Cre^*-induced *Atf4* inactivation, there is a clear separation of control and mutant β-cells but not other cell types in an UMAP (Fig. 5B), suggesting that the gene expression changes in islets largely lie within β-cells.

**Fig. 5.**
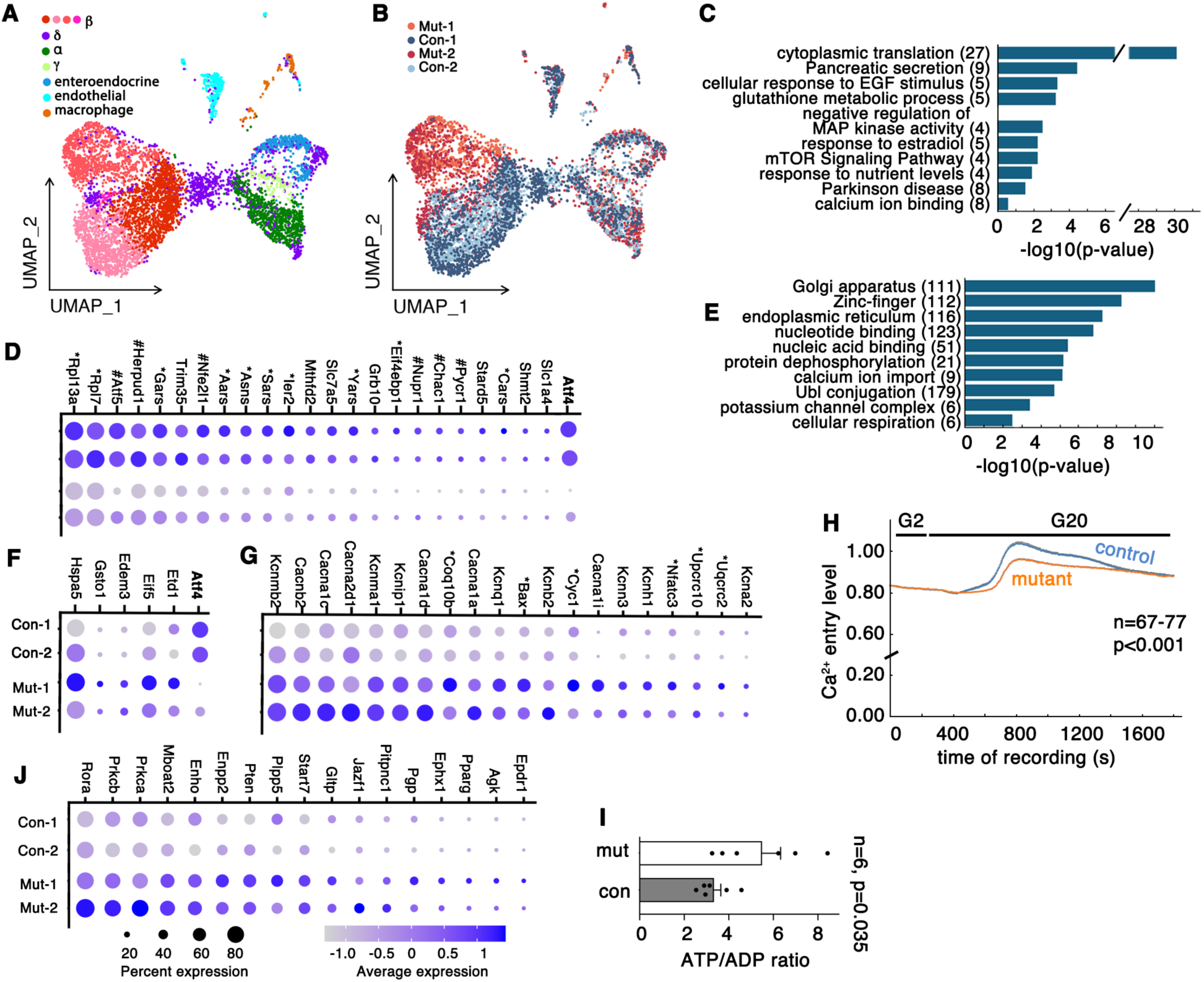
ScRNA-seq identifies DEGs between control and Atf4-deficient β-cells that correlate with functional alterations of mutant β-cells. Islets from 2-months old mice were handpicked and dissociate into single cells for scRNA-seq. Two controls (Con-1 and Con-2, *Atf4^F/F^*) and two mutant (Mut-1 and Mut-2, *Atf4^F/F^; Ins1^Cre^*) samples were used. (A) An UMAP with all identified cells that have high quality transcriptomes. The β-cells were presented as 4 subpopulations, a heterogeneity shown by others as well (but not further followed in this study). (B) An UMAP highlighting the cell types with shifted gene expression according to genotypes. (C) The top 10 down-regulated pathways in *Atf4*-deficient β-cells. The numbers in parenthesis are the DEGs belonging to each pathway. (D) Dot plots of DEGs that have Atf4 binding sites within 3-kb of their TSS. *: genes involved in translation. #: genes involved in stress response. Note that *Atf4* expression was included (far right) as control to show the degree of Atf4 deletion in β-cells. (E) The top 10 up-regulated pathways in *Atf4*-deficient β-cells, with the number of genes involved noted. (F, G) Dot plot of several upregulated genes which are putative Atf4 targets (F) or non-targets (G). In F, only genes involved in translation and stress response are shown. In G, only channel and respiratory genes (marked with *) are shown. (H) Glucose-induced Ca^2+^ in four months-ol-d islets, assayed by FURA2AM, quantified with F340/F380 ratios. Shown are mean + SEM. P value is from two-way ANOVA, repeated measure. (I) ATP/ADP ratios of control and mutant islets. (J) Dot plots of several DEGs involved in lipid metabolism. Note that in all dot plots, percent expression refers to the portions of cells with detectable messages. Average expression is mRNA reads averaged against all cells and then log transformed.

The differential analysis identified 1,161 DEGs between mutant and control β-cells, with 161 down- and 1,000 up-regulated in mutant β-cells (Table S1). Functional Clustering assays with DAVID showed that the most enriched biological process in the down-regulated DEGs is cytoplasmic translation, with 27 of the 161 genes coding for ribosomal proteins (Fig. 5C, Table S1). Biological processes such as pancreatic secretion, glutathione metabolic process, and response to nutrient levels are also enriched, likely related to compromised function and/or oxidative stress response in mutant cells (Fig. 5C). Taking advantage of the known 415 Atf4-binding sites in fibroblasts (26), we found putative Atf4 binding sites within three kilobase of transcription starting sites (TSS) in 23 of the 161 genes (Fig. 5D), representing a ∼6.0-fold enrichment over random overlapping (*p*=5.5E-12, hypergeometric assays). Notably, 10 of these 23 genes are involved in translational initiation or aa-tRNA production and six in stress response (Fig. 5D. marked with * or #, respectively). These findings highlight the roles of Atf4 in directly activating protein translation and stress response. Gene Set Enrichment Analysis (GSEA) revealed similar but more detailed GO and KEGG pathways from these DEGs, presented in Table S2.

The biological processes that are upregulated in the *Atf4* mutant β cells include Golgi, ER, channels (i. e. K^+^ and Ca^2+^ channels), and respiration (Fig. 5E). These overall findings, combined with the secretory behaviors of Atf4-deficient β cells, highlight the roles of Atf4 in depressing vesicular production/trafficking and stress response. Notably, 25 of the 1,000 genes are known Atf4 targets (Table S1) (26). This small number of genes *per se* represents no significant enrichment over random overlapping between Atf4 targets and upregulated genes (*p*=0.43). Yet two of the 25 genes are involved in translational repression (*Etf1* and *Eif5*) (47,48), while three [*Edem3*, *Gsto1* (encoding glutathione transferase omega 1), and *Hspa5*] in stress response (17,49,50)(Fig. 5F). These latter results are consistent with Atf4 acting as both transcriptional activator and repressor, both participating translational regulation and stress response. They also suggest that most of the upregulated genes are activated in response to the lack of Atf4 function instead of being directly repressed by Atf4.

### Loss-of *Atf4* compromises glucose-induced Ca^2+^ influx but not mitochondrial function

We next explored if the altered gene expression in *Atf4* mutant β-cells can explain some of the mutant β-cell behaviors. For this goal, we first examined the glucose-induced Ca^2+^ influx, because two of the highly upregulated pathways are related to Ca^2+^ (*Cacna1d, Cacna1a, Cacna1c,* and *Cacna1i*) and potassium channels (*Kcnka2, Kcnb2, Kcnh1, Kcnma1, Kcnmb2, Kcnip1, Kcnq1,* and *Kcnn3*) (Fig. 5F). The hypothesis is that the combined activities of these factors can shift the glucose-induced Ca^2+^ influx. Four-months old islets, after being cultured at G11 for two days have significantly reduced Ca^2+^ influx in response to G20 (Fig. 5H), corresponding to the reduced GSIS in mutant islets under this condition (Fig. 3E). This reduced Ca^2+^ influx does not seem to be caused by reduced ATP production, because the mutant β-cells have higher expression of several genes in cellular respiration (including *Bax*, *Coq10b*, *Cyc1*, *Nfatc3*, *Uqcrc2*, and *Uqcrc10*) (Fig. 5G. Table S1). More importantly, the mutant islets have higher levels of ATP/ADP ratio when stimulated by G20 (Fig. 5I). These combined results suggest that the combined upregulation of Ca^2+^ and K^+^ channel proteins have altered the electrical properties of mutant β-cell membrane, resulting in lowered overall Ca^2+^ influx.

We last examined if any DEG is potentially involved in protecting free fatty acid-induced β-cell dysfunction. We focused on genes involved in lipid metabolism which may regulate the accumulation of specific metabolites that causes lipotoxicity (10). Manual gene-by-gene examination of all up-regulated DEGs, combined with DAVID analysis, identified 17 lipid-metabolism and signaling-related genes (Fig. 5J). These include *Jaz1, Pparg, Pten, Prkca,* and *Prkcb* that is involved in general lipid metabolism and signaling. These also include *Plpp5, Mboat2, Agk,* and *Enpp2* that is related to phospholipid metabolism. The increased *Agk* (encoding an acyl-glycerol kinase) expression in *Atf4*-deficient β-cells is particularly intriguing, because this gene was shown to not only regulate lipid metabolism, but also mitochondrial function (51), the primary target of lipotoxicity.

## Discussion

Here, we show that aged *Atf4-*deficeint β-cells are sensitive to acute hyperglycemia-induced dysfunction but are resistant to hyperlipidemia-induced failure. Corresponding to these different responses, *Atf4*-deficient β-cells have changed expression of several sets of genes that regulate different aspects of β-cell function and stress response. These results highlight the dilemma that β-cells face under glucolipotoxicity. They activate Atf4 to alter the transcriptome to handle high glucose-induced stress but this change exacerbates their free fatty acid-induced dysfunction. We propose this as a major mechanism underlying the synergizing effect of glucolipotoxicity on β-cell failure. Thus, manipulating Atf4-regulated genes that are involved in high glucose or high free fatty acid-related process may preserve functional β-cell mass to delay/avoid T2D development.

Obesity is usually associated with hyperglycemia and hyperlipidemia. The former triggers cascades of changes in insulin secretion/biosynthesis, gene expression, and cellular stress (1,52). The latter can directly enhance insulin secretion while also increasing the levels of metabolites such as ceramides that induce cellular stress (10,53). Thus, proper stress response is an essential process that maintains β-cell homeostasis and function. However, overactivation of stress response can lead to β-cell failure as well, underscoring the importance of understanding their regulation (1,14,17,54).

Atf4 is a key mediator of stress response with context dependent roles (26,55). We have shown that its overexpression compromises β-cell identity in mice (15). Two recent studies showed that *Atf4* in β-cells is required for glucose tolerance under normal feeding conditions and required for β-cell identity/function under ER stress (28). A caveat is that *Atf4* inactivation was achieved using *Rip^Cre-TG^*, a transgene that compromises β-cell function (29) but not included as controls (22,28). Furthermore, the Atf4-dependent gene expression in β-cells is not known. By using a non-toxic Cre line, we show that *Atf4* is dispensable for β-cell function under normal physiology, but it is needed under high glucose-related stress, with aging exacerbate mutant β-cell dysfunction. More importantly, we uncover the opposing roles of Atf4 in β-cells when facing acute high glucose or high free fatty acid treatment, at the levels of both secretory function and identity-gene expression. These results can explain not only the synergistic effects of hyperglycemia and hyperlipidemia on β-cell failure, but also the lack of convincing evidence of lipotoxicity as a cause for β-cell failure (56).

How Atf4 regulates the differential β-cell response to high levels of glucose or free fatty acids is currently not clear. Our β-cell specific gene expression analysis suggests several possibilities. Specifically, we found that the Atf4-deficient β-cells have reduced pathways involved in translation, secretion, and glutathione metabolism (Fig. 5C), which likely sensitize the mutant cells to high-glucose induced failure. Along a similar line, we detected increased expression of several genes involved in lipid metabolism/signaling in *Atf4*-deficient β-cells, including *Agk*, an enzyme that facilitate phospholipid production and mitochondrial function (51). These gene products may shift the flux of lipid metabolites and/or signals, leading to protected β-cell identity and function via mitochondria-related pathways. In addition, the upregulated Atf4 target genes in Atf4-deficient β-cells have several that participate translational repression and stress response (Fig. 5F), with *Gsto1* shown to protect cells from oxidative stress (57-59). Thus, these latter genes, individually or cooperatively, may protect Atf4-deficient β-cells from high faty acid-induced failure. Future studies to examine these possibilities will be extremely informative.

We cannot explain why α-cells in mutant mice have higher levels of proliferation. It is possible that paracrine signals from the *Atf4*-deficient β-cells are involved. Alternatively, these mutant mice may induce α-cell proliferation via systematic signals. Lastly, we do not know the reason for the discrepancy between ours and others’ findings when *Rip^Cre-TG^* was used for *Atf4* deletion. It is possible the phenotypes could be attributed to the hGH overexpression in the *Rip^Cre-TG^* mice. It is also possible that this transgene has been partially silenced in our facility, resulting in Atf4 inactivation in only a portion of β-cells. Thus, we propose caution when using *Rip^Cre-TG^*, when these mice should be included in all studies as controls and its Cre activity be monitored using reporter-activation or loss-of targeted gene products.

## Author contributions

MY, SN, and RH, did IF assays and performed GSIS tests. SN, YX, and RH produced/genotyped mice and performed IPGTT. RH also performed ATP/ADP assays. YZ and QL analyzed scRNA-seq data. YX, AJS, KL performed scRNA-seq. PD and DJ did Ca^2+^ recording. CA, RS, and GG conceptualized the study. All authors participated manuscript writing/proofing.

## Guarantor Statement

GG is guarantor of the study, with full access to and takes responsibility for the integrity of the data and the accuracy of the data analysis.

## Previous presentation information

The work has not been presented anywhere.

## Description of supplemental materials

### Supplemental tables

**Table S1. DEGs between 2-months-old control and Atf4-deficient β-cells. Related to Fig. 5**. Average expression in two controls and two mutants are presented. Only genes with an adjusted *p* value below 0.05 were listed. Note that genes with Atf4-binding sites within 3-kb 5’ to TSS were also marked.

**Table S2. Lists of GO terms and KEGG pathways that are enriched in control or mutant β-cells.** The results were derived using GSEA, complementing the functional clustering using DAVID.

### Supplemental Figures

**Fig. S1.**
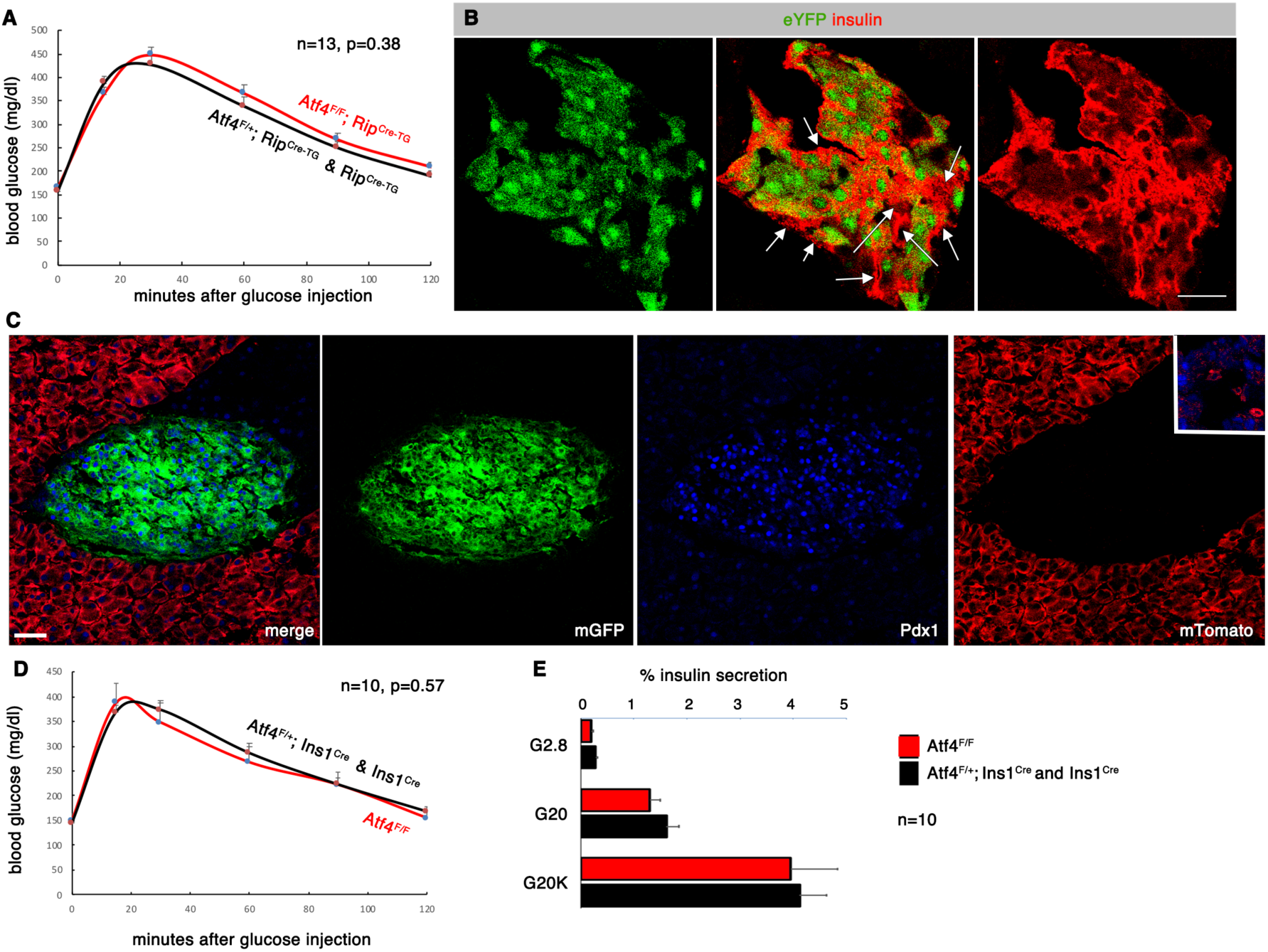
Evaluating *Rip^Cre-TG^* and *Ins1^Cre^*-mediated *Atf4* inactivation in β-cells. (A) IPGTT in 6-months-old mutant (*Atf4^F/F^; Rip^Cre-TG^*) and control (*Rip^Cre-TG^* and *Atf4^+/F^; Rip^Cre-TG^*) mice. Both male and females are included. (B) Evaluation of detectable Cre in *Rip^Cre-TG^* mice, using a R26R^eYFP^ reporter. White arrows point at several β-cells without detectable Cre. (C) Evaluation of detectable Cre in *Ins1^Cre^* mice, using a MT/MG reporter (express mTomato without detectable Cre but GFP with Cre). Note that Pdx1 staining is used here to locate islets. Inset in the mTomato channel show three cells within islets that do not express Pdx1 and active Cre. (D) IPGTT in 6-months-old *Atf4^F/F^* and *Atf4^+/F^; Ins1^Cre^* plus *Ins1^Cre^* mice, both males and females included. (E) GSIS from 6-months-old *Atf4^F/F^* and *Atf4^+/F^; Ins1^Cre^* plus *Ins1^Cre^* mice (from both male and female mice). Note that in A, D, E, mean + SEM were shown. P values were calculated with t-test, two-tailed type 2 error. Bars = 20 μm. Sections in B and C are both from male mice, similar to that of tested females.

**Fig. S2.**
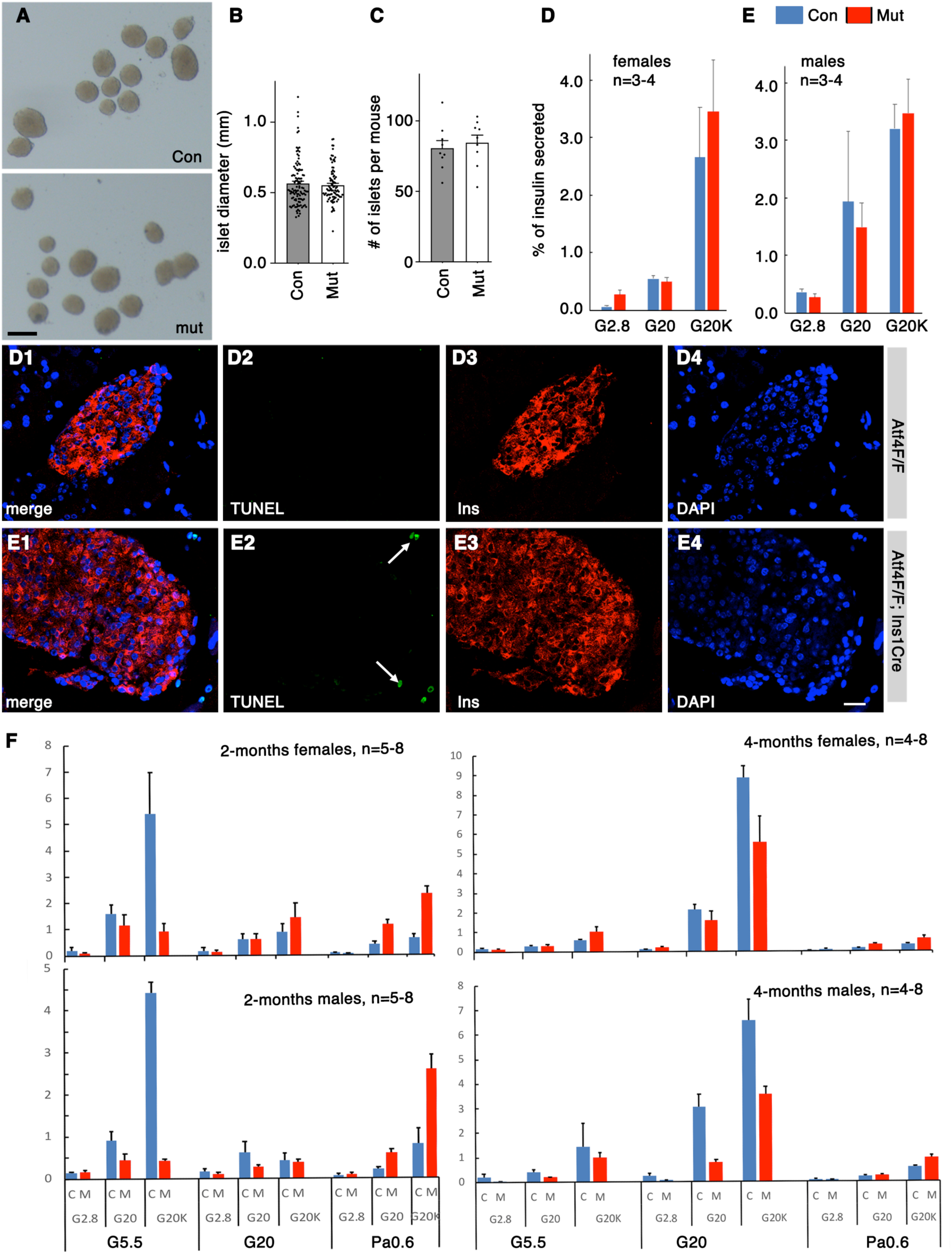
Characterization of several islet properties in control (Con or *Atf4^F/F^*) and mutant (Mut or *Atf4^F/F^; Ins1^Cre^*) mice. (A) Examples of islets from 1-month-old male mice. Scale bar, 100 μm. (B, C) Average islet size and islet numbers per mouse (1-month old, both males and females are included, with results combined). (D, E) Insulin secretion from isolated islets of 1-month-old mice, with male and female data presented separately. (F, G) TUNEL assays of control and mutant that have induced hyperglycemia with S961 infusion. Note arrows in G2 point at a couple of TUNEL signal positive cells outside of islets, showing the cell death of none β-cells. Scale bars, 20 μm. Used are sections from males, with females having identical results. (H) GSIS from 2- and 4-months-old male and female mice that were treated with G5.5, G20, and Pa0.6 for 48 hours *in vitro* before secretion assays. In F, “C”=control, “M” mutants. Note that in B-E and F, mean + SEM were shown. P values were calculated with t-test, two-tailed type 2 error.

**Fig. S3.**
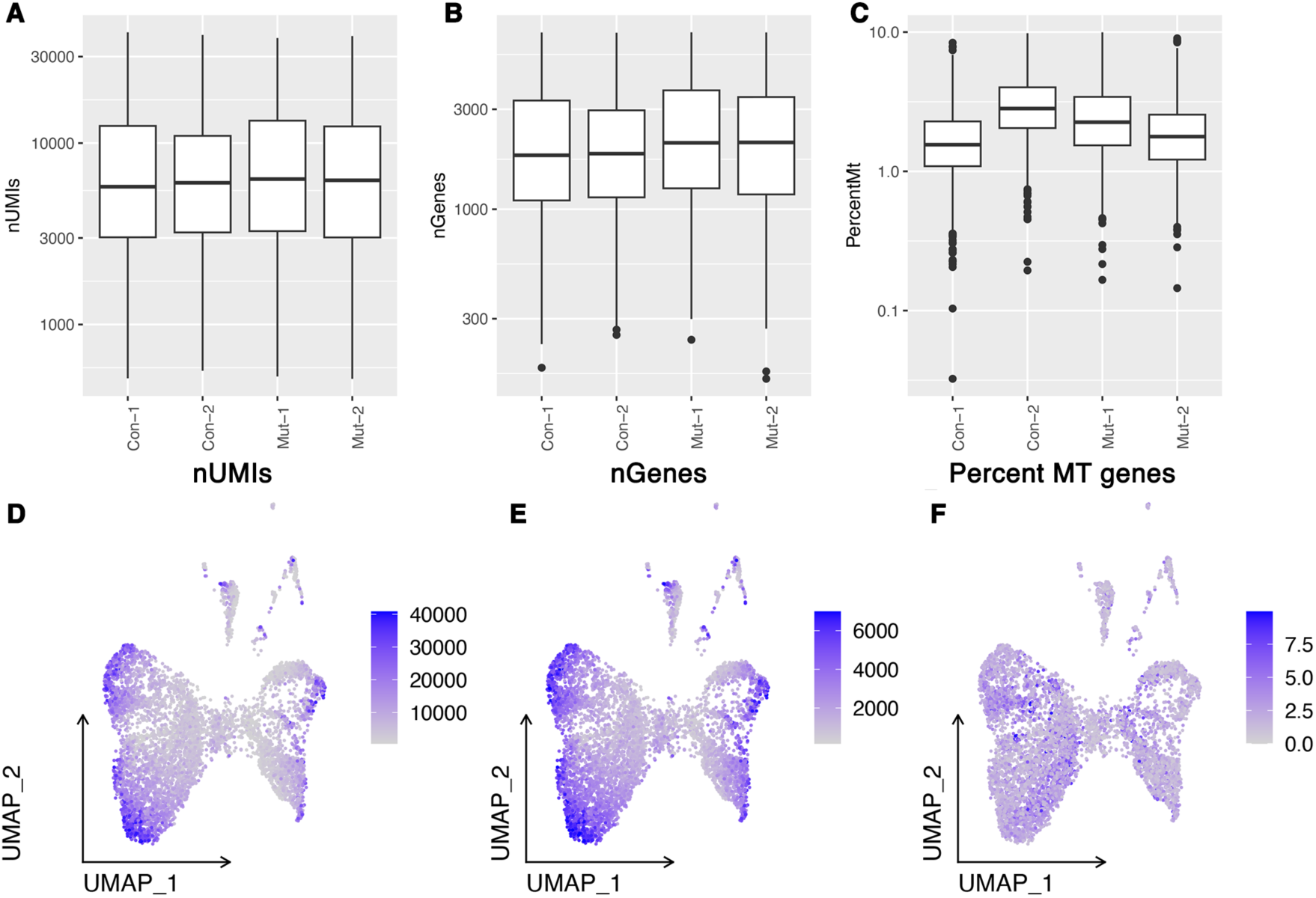
Quality control of scRNAseq data. (A) Number of UMIs per cell for each sample. (B) Number of genes per cell for each sample. (C) Percentage of reads mapped to mitochondrial genes per sample. (D) Distribution of number of UMIs on UMAP. (E) Distribution of number of genes on UMAP. (F) Distribution of percentage of mt-RNAs on UMAP.

